# Evidence for Normalization as a Fundamental Operation Across the Human Visual Cortex

**DOI:** 10.1101/2021.05.21.445228

**Authors:** Narges Doostani, Gholam-Ali Hossein-Zadeh, Maryam Vaziri-Pashkam

## Abstract

Divisive normalization of the neural responses by the activity of the neighboring neurons has been proposed as a fundamental operation in the nervous system based on its success in predicting neural responses recorded in primate electrophysiology studies. Nevertheless, experimental evidence for the existence of this operation in the human brain is still scant. Here, using functional MRI, we explored the role of normalization across the visual hierarchy in the human visual cortex. Using stimuli form the two categories of human bodies and houses, we presented objects in isolation or in clutter and asked participants to attend or ignore the stimuli.Focusing on the primary visual area V1, the object-selective regions LO and pFs, and the category-selective regions EBA and PPA, we first modeled single-voxel responses using a weighted sum, a weighted average, and a normalization model and demonstrated that the response to multiple stimuli could best be described by a model that takes normalization into account. We then explored the observed effects of attention on cortical responses and demonstrated that these effects were predicted by the normalization model, but not by the weighted sum or the weighted average models. Our results thus provide compelling evidence for normalization as a canonical computation operating in the human brain.

## Introduction

The brain makes use of fundamental operations to perform neural computations in various modalities and different regions. Divisive normalization has been proposed as one of these fundamental operations. Under this computation, the response of a neuron is determined based on its excitatory input divided by a factor representing the activity of a pool of nearby neurons ^1,2,3^. Normalization was first introduced based on responses in the cat primary visual cortex ^1^, and evidence of its operation in higher regions of the monkey visual cortex has also been demonstrated both during passive viewing ^4^ and when attention is directed towards a stimulus ^5,6,7,8^.

Normalization has also been proposed as a critical operation in the human brain based on evidence demonstrating sub-linear addition of responses to multiple stimuli in the visual cortex ^9^. Nevertheless, in lieu of directly testing the normalization model to resolve multiple-stimulus representation, several previous studies have shown that a weighted average model can account for multiple-stimulus responses in the monkey ^10^ or human ^11,12,13^brain. The only exception is a recent electrophysiology study, which showed that in the category-selective regions of the monkey brain, a winner-take-all, but not averaging, rule can explain neural responses in many cases ^4^. Bao and Tsao ^4^ further demonstrated that the normalization model predicts such winner-take-all behaviour. It is not clear whether this discrepancy has emerged as a result of different explored regions of the brain, or due to the diversity in stimuli or the task performed by the subjects.

In addition to regional computations for multiple-stimulus representation, the visual cortex relies on top-down mechanisms such as attention to select the most relevant stimulus for detailed process-ing ^14,15,16^. Attention works through increasing the response gain ^17,18^ or contrast gain ^19,20^ of the attended stimulus. Previous studies have shown the role of normalization in regulating the gain of the attended stimulus in the monkey brain ^7,8,12,21^. Normalization has also been speculated to underlie response modulations in the human visual cortex in the presence of attention^9^. However, no study to date has directly tested the validity of the normalization model in predicting human cortical responses ^9,13,22^.

To fill this gap and to explore the discrepancies reported about multiple-stimulus responses, here, we aimed to evaluate the predictions of the normalization model against observed responses to visual objects in several regions of the human brain in the presence and absence of attention. In an fMRI experiment using conditions with isolated and cluttered stimuli and recording the response with or without attention, we provide a comprehensive account of normalization in different regions of the visual cortex, showing its success in adjusting the gain of each stimulus when it is attended or ignored. We also demonstrate that normalization is in fact close to averaging in the absence of attention, as previously reported by several studies ^10,11,13^, but that the results of the two models diverge in the presence of attention. Our work in the human brain, along with previous studies of normalization in the monkey brain, provides compelling evidence for normalization as a canonical computation in the primate brain.

## Results

### Attention modulates responses to isolated and paired stimuli

In a blocked-design fMRI paradigm, human participants (n=19) viewed semi-transparent gray-scale stimuli from the two categories of houses and human bodies (Figure 1a). Each experimental run consisted of one-stimulus (isolated) and two-stimulus (paired) blocks, with attention directed either to one of the two objects or to the color of the fixation point. The experiment, therefore, had a total number of 7 conditions (four isolated and three paired conditions, see Figure 1c). In paired blocks, we superimposed the two stimuli to minimize the effect of spatial attention and force participants to use object-based attention (Figure 1b,c). Independent localizer runs were used to localize the primary visual cortex (V1), the object-selective regions in the lateral occipital cortex (LO) and posterior fusiform gyrus (pFs), the extrastriate body area (EBA), and the parahippocampal place area (PPA) for each participant (Figure 1d).

**Figure 1:**
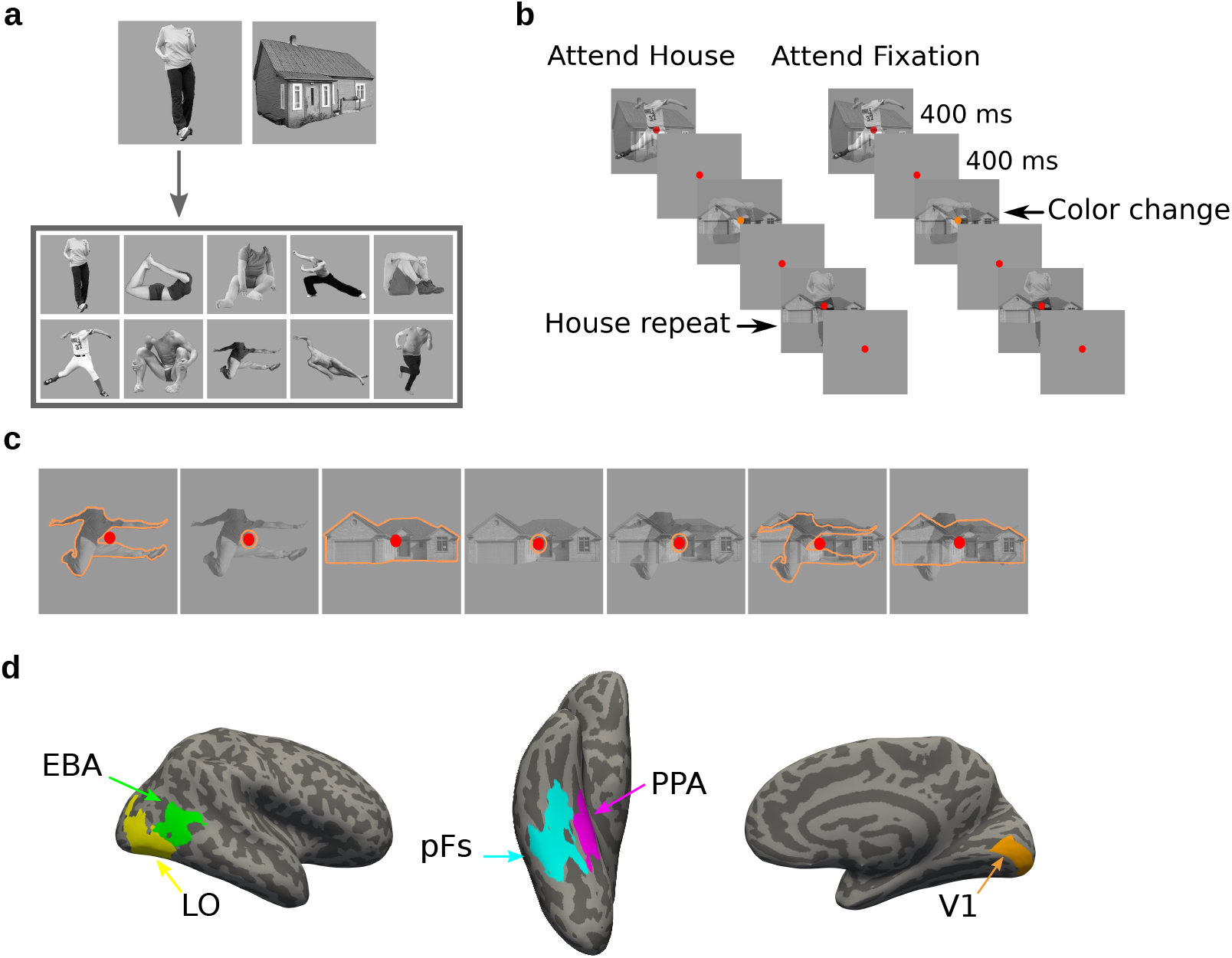
Stimuli, paradigm and regions of interest. (a) The two stimulus categories (body and house), with the ten exemplars of the body category. (b) Experimental paradigm including the timing of the trials and the inter-stimulus interval. Inthe example block depicted on the left, both stimulus categories were presented, and the participant was cued to attend to the house category. The two stimuli weresuperimposed in each trial, and the participant had to respond when the same stimulus from the house category appeared in two successive trials. The color of the fixation point randomly changed in some trials from red to orange, but the participants were asked to ignore the color change. The example block depicted on the rightillustrates the condition in which stimuli were ignored and viewers were asked to attend to the fixation point color, and respond when they detected a color change.(c) The 7 task conditions in each experimental run. For illustration purposes, we have shown the attended category in each block by orange outlines. The outlineswerenot present in the actual experiment. (d) Regions of interest for an example participant, including the primary visual cortex V1, the object-selective regions LO and pFs, the body-selective region EBA, and the scene-selective region PPA.

To examine the response variation in different task conditions, we fit a general linear model and estimated the coefficients for each voxel in each task condition. We then defined the preferred (P) and null (N) stimulus categories for each voxel in the five regions of interest (ROIs), including V1, LO, pFs, EBA, and PPA, according to the voxel’s response to isolated body and isolated house conditions. We then rearranged the seven task conditions according to each voxel’s preferences. The conditions are hereafter referred to as: P^at^, P^at^N, PN^at^, N^at^, P, PN, N, with P and N denoting the presence of the preferred and null stimuli, respectively, and the superscript *at* denoting the attended category. Mean voxel responses in the five ROIs for all task conditions are illustrated by navy lines in Figure 2a,b. The change in the BOLD response across task conditions demonstrates how presented stimuli and attention affect voxel responses.

**Figure 2:**
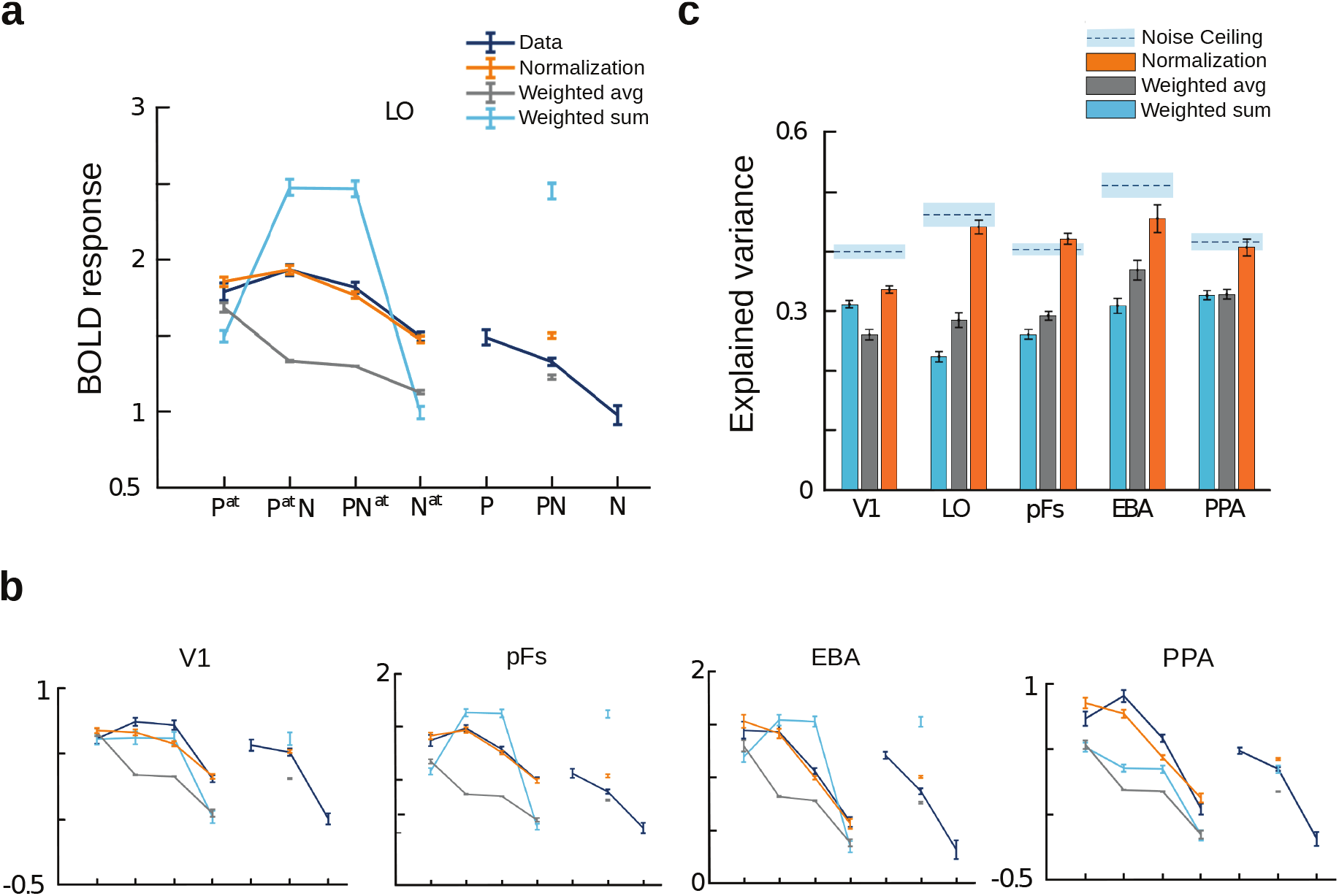
Divisive normalization explains voxel responses in different stimulus conditions. (a-b) Average fMRI responses and model predictions in the five regions of interest. Navy lines represent average responses. Light blue, gray, and orange lines show the predictions of the weighted sum, the weighted average, and the normalization models, respectively. The x-axis labels represent the 7 task conditions, P^at^, P^at^N, PN^at^, N^at^, P, PN, N, with P and N denoting the presence of the preferred and null stimuli and the superscript *at* denoting the attended category. For instance, P refers to the condition in which the unattended preferred stimulus was presented in isolation, and P^at^N refers to the paired condition with the attended preferred and unattended null stimuli. Error bars represent standard errors of mean. (c) Mean explained variance, averaged over voxels in each region of interest for the 5 conditions predicted by the three models. Light blue, gray, and orange bars show the average variance explained by the weighted sum, weighted average, and normalization models, respectively. Error bars represent the standard errors of mean. Dashed lines above each set of bars indicate the noise ceiling in each ROI, with the gray shaded area representing the standard errors of mean.

### Divisive normalization explains voxel responses in different stimulus condi-tions

We used the three models of weighted sum, weighted average, and normalization to predict voxel responses in different task conditions. The weighted sum model which has the advantage of simplicity is a linear model according to which the response to multiple stimuli is determined by the sum of the responses to each individual stimulus presented in isolation, and attention to each stimulus increases the part of the response associated with the attended stimulus:

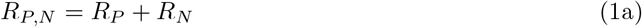

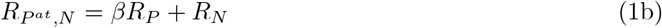

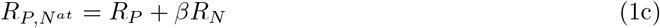

Here, R_P, N_ denotes the response elicited with the preferred and null stimuli present in the receptive field, and R_P_and R_N_ denote the response to isolated preferred and null stimuli, respectively. The superscript *at* specifies the attended stimulus, and the stimulus is ignored otherwise. The response related to the attended stimulus is weighted by β, which is the parameter related to attention.

The weighted average model has previously been successful in explaining responses to multiple stimuli in the absence of attention ^10,11,23^. According to this model, the response to paired stimuli is the average of the response to each stimulus presented alone, weighted by β as the attention-related parameter:

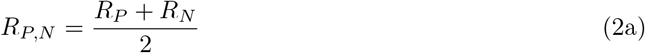

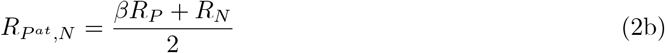

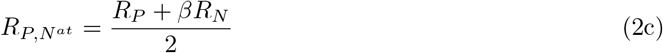

The normalization model of attention is a more elaborate model based on which each neuron receives an excitation caused by a stimulus present in its receptive field, in addition to a suppression due to that stimulus because the neighboring neurons are also excited by the same stimulus. This model can thus be described using divisive normalization with a saturation term in the denominator ^1,2,5,7^:

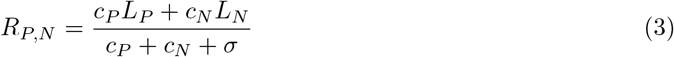

Here, L_P_ and L_N_ denote the excitatory drive induced by the preferred or the null stimulus, respectively and σ represents the semi-saturation constant. c_P_and c_N_ are the respective contrasts of the preferred and null stimuli. Zero contrast for a stimulus denotes that the stimulus is not present in the visual field. When attention is directed towards one of the stimuli, both the excitation and the suppression related to that stimulus are enhanced. We can therefore rewrite Equation 3 as:

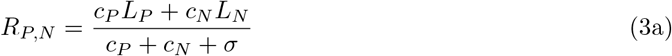

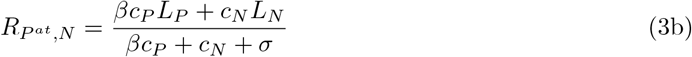

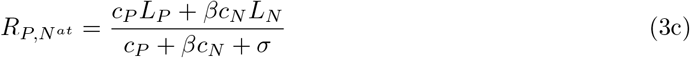

To compare these models, we split the fMRI data into two halves (odd and even runs) and estimated the model parameters separately for each voxel of each participant using the first half of the data. All comparisons of data with model predictions were made using the second left-out half of the data. Note that this independent prediction is critical since the numbers of parameters in the three models are different. Possible over-fitting in the normalization model with more parameters will not affect the independent predictions. Figure 2a,b shows average BOLD responses for the five modeled task conditions in all ROIs (the two isolated ignored conditions P and N were excluded as they were used to predict responses in the remaining five conditions, see supplementary methods). As evident in the figure, the predictions of the normalization model (in orange) very closely follow the observed data (in navy), while the predictions of the weighted sum and weighted average models (light blue and gray, respectively) have significant deviations from the data. To quantify this observation, the goodness of fit was calculated for each voxel by taking the square of the correlation coefficient between the predicted model response and the respective fMRI responses across the five modeled conditions. Noise ceiling was calculated by taking the r-squared of the correlation between voxel responses in the two halves of data. As illustrated in Figure 2c, the normalization model (orange bars) explained the variance in the responses significantly better than the weighted average and the weighted sum models in all regions of interest. Moreover, the explained variance by the normalization model predictions was not significantly different from the noise ceiling in LO, pFs and PPA (*ts* < 1.7, *ps* > 0.11).

Interestingly, just focusing on the paired condition in which none of the stimuli were attended (the PN condition), the normalization model was not necessarily better than the weighted average model (the gray and orange isolated data points on the subplots of Figure 2 are similarly close to the navy line). For this condition, the normalization model was better than the weighted average model in V1 and PPA (*ts* > 2.7, *ps* < 0.013) but the two models showed no difference in predicting the data in all other regions (*ts* < 1.7, *ps* > 0.11). These results are in line with previous studies which have shown that the weighted average model can predict neural and voxel responses in the absence of attention ^10,11,13^. However, here we show that in the presence of attention, the normalization model has a significant advantage.

### Normalization accounts for the change in response with the shift of attention

Next, comparing the responses in different conditions, we observed two features in the data. First, for the paired conditions, shifting attention from the preferred to the null stimulus caused a reduction in voxel responses. We calculated this reduction in response for each voxel by (*P^at^N — PN^at^*) (Figure 3a, top panel). This response change was significantly greater than zero in all ROIs (*ts* > 5.9, *ps* < 1.5 × 10^−5^) except V1 (t(18) = 0.18, p = 0.43). Because the same stimuli were presented in both conditions but the attentional target changed from one category to the other, this change in response could only be related to the shift in attention and the stimulus preference of the voxels.

**Figure 3:**
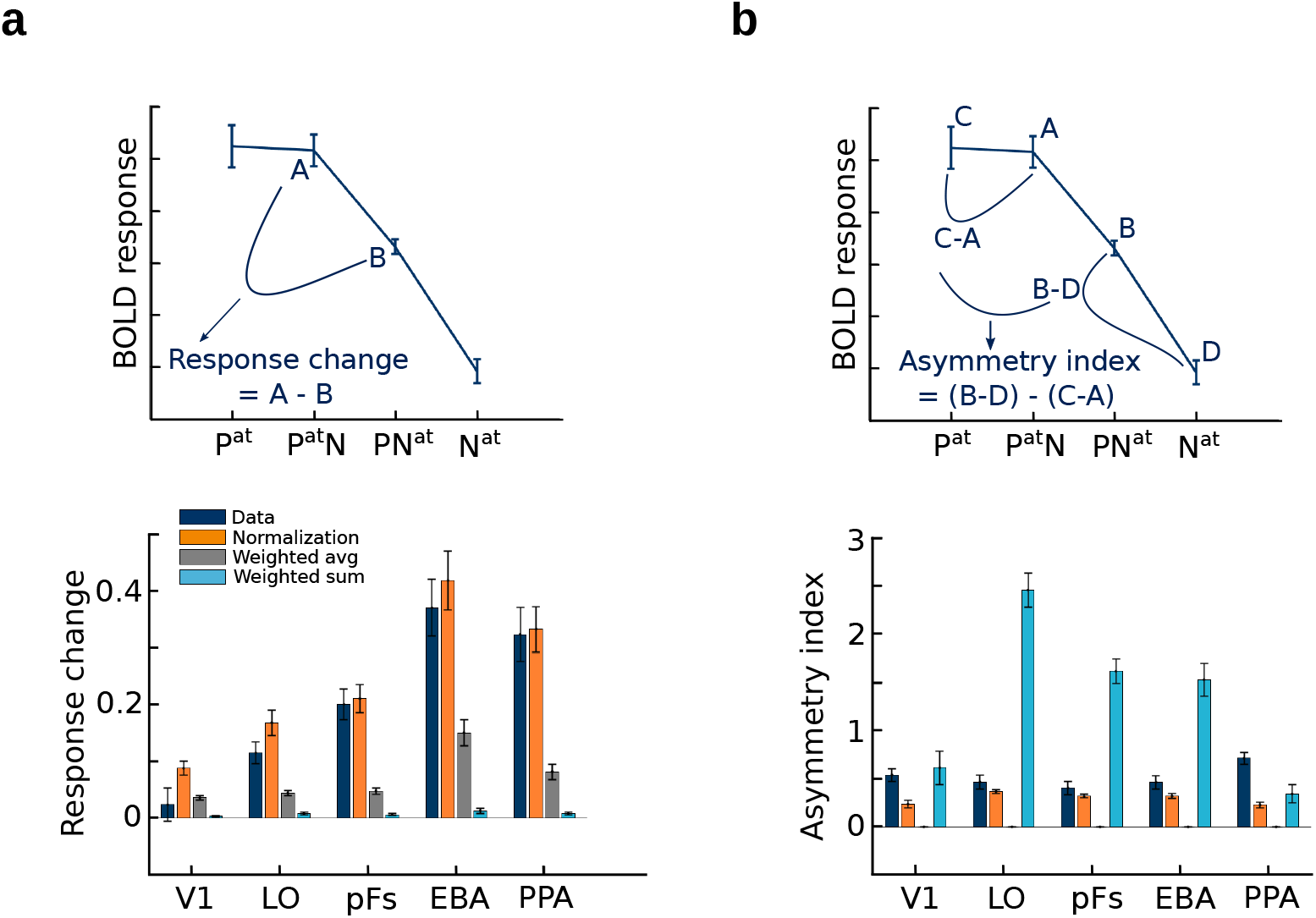
Normalization accounts for the observed effects of attention. (a) Top: Change in BOLD response when attention shifts from the preferred to the null stimulus in the presence of two stimuli, illustrated here for EBA. Bottom: The observed response change and the corresponding amount predicted by different models in different regions, calculated as illustrated in plot A. Error bars represent the standard errors of mean. (b) Top: The observed asymmetry in attentional modulation for attending to the preferred versus the null stimulus, depicted for EBA. Bottom: The observed and predicted asymmetries in attentional modulation in different regions, calculated as illustrated in plot B. Error bars represent the standard errors of mean.

We then compared this observed feature of the data with what was predicted by the three models to investigate model results in more detail. As illustrated in the bottom panel of Figure 3a, the orange bars depicting the predictions of the normalization model were very close to the navy bars depicting the observations in all ROIs, while the predictions of the weighted sum and weighted average models (light blue and gray bars, respectively) were significantly different from the data. The normalization model therefore predicted that shifting attention from the preferred to the null stimulus in the paired conditions reduces voxel responses. The amount of this reduction in response was also the same as what we observed in the data in all regions (*ts* < 1.94, *ps* > 0.068) except for LO (t(18) = 2.87, p = 0.01). The weighted average model predicted very little response reduction, significantly different from what was observed in the data in all regions (*ts* > 4.2, *ps* < 0.0005) except for V1 (t(18) = 0.32, p = 0. 75). The weighted sum model predicted no significant reduction in response.

### Asymmetry in attentional modulation is explained by the normalization model

The second feature we observed was that the effect of the unattended stimulus on the response depended on voxel selectivity for that stimulus, with the unattended preferred stimulus having larger effects on the response than the unattended null stimulus. Attending to the preferred stimulus in the presence of the null stimulus caused the response to approach the response elicited when attending to the isolated preferred stimulus. Therefore, attention effectively removed the effect of the null stimulus. However, attending to the null stimulus in the presence of the preferred stimulus did not eliminate the effect of the preferred stimulus and yielded a response well above the response elicited by attending to the isolated null stimulus. This is the first time such asymmetry has been reported in human fMRI studies, but similar results have been found in monkey electrophysiology studies ^6,7,14^. To quantify the observed asymmetry we calculated an asymmetry index for each voxel by (*PN^at^ – N^at^*) —(P^at^ – P^at^N) which is illustrated in the top panel of Figure 3b. This index was significantly greater than zero in all regions (*ts* > 6, *ps* < 10^−5^).

Here too, the normalization model was better at predicting the observed asymmetry in the data. The bottom panel of Figure 3b illustrates the asymmetry indices for the data and the three models in all regions. The normalization model predicted the asymmetry index more accurately than the weighted sum model in all regions (ts > 5.7, ps < 10^−5^) except for V1 and PPA (ts <1.9, ps > 0.07). Notably, the predicted index by the normalization model in LO and pFS was not significantly different from the observed index (ts < 1.7, ps > 0.11). The weighted average model predicted no significant asymmetry in attentional modulation.

## Discussion

Here, modeling responses to single voxels, we explored the validity of the normalization model in predicting voxel responses in the human visual cortex. While several electrophysiology studies have explored normalization in the monkey brain^4,7,8,21^, and although normalization has also been proposed to operate in the human brain ^9,22^, no study to date has directly tested its validity in the human brain. Here, for the first time, we demonstrated the success of the normalization model and its superiority to the weighted sum and weighted average models in predicting cortical responses to single and multiple stimuli. Next, we investigated the effect of attention on multiple-object representation. Defining preferred and null stimuli for each voxel, we showed that when attention shifts from the preferred to the null stimulus, there is a significant reduction in response in object-selective and category-selective regions of interest, but not in the primary visual area. Furthermore, this response reduction increases as we move to higher regions across the hierarchy, consistent with speculations of greater effects of top-down attention on higher regions of the visual cortex ^24^. We demonstrated that the normalization model, but not the weighted sum or the weighted average models, could account for this response reduction observed in different regions. Finally, our results indicated an asymmetry in attentional modulation when attending to the preferred versus the null stimulus. We demonstrated that while attention to a preferred stimulus almost eliminates the effect of the ignored null stimulus, attention to the null stimulus does not remove the effect of the preferred stimulus. Unlike response change with the shift of attention, which increases across the hierarchy, the asymmetry measured by our defined index stays approximately the same in all regions, indicating its dependence not on the top-down attentional signal but on the normalization computation performed in each region.

Normalization has been reported to cause neural populations to operate in the two regimes of averaging and winner-take-all based on stimulus contrasts ^25^. Here, we showed that in the presence of attention, responses can deviate from the averaging regime even without change in contrast. We observed a winner-take-all behavior when the preferred stimulus was attended since its higher response along with its increase in gain due to attention made it a much stronger input compared to the ignored null stimulus. On the other hand, when the null stimulus was attended, the response was closer to an average than a max-pooling response. This result explains why several previous studies in the object-selective cortex indicated averaging as the rule for multiple-stimulus representation ^10,11^, while studies in category-selective areas reported a winner-take-all mechanism ^4^. We therefore extend previous reports of multiple-stimulus representation by demonstrating that the response is not only related to stimulus contrast ^25^ or neural selectivity ^4^, but that a combination of bottom-up and top-down signals act to yield a response that can be the average of the isolated responses, or a winner-take-all response, or somewhere between the two. We also demonstrate for the first time that the normalization model is superior to the weighted average model, which has often been used in lieu of the normalization model ^10,11,13^, in its ability to account for fMRI responses in the presence of attention.

It is noteworthy that here, we are looking at the BOLD responses. We are aware of the limitations of the fMRI technique as the BOLD response is an indirect measure of the activity of neural populations. We hope that future research will directly test the effectiveness of the normalization model in predicting neural responses in the human brain.

In sum, our results indicate that the normalization model can explain responses at the voxel level beyond the primary visual cortex and across the visual hierarchy, especially in category-selective areas of the human visual cortex, with and without attention, and in conditions with isolated or cluttered stimuli. We therefore provide strong evidence for divisive normalization as a canonical computation operating in the human brain.

## Methods

### Participants

A total of 21 healthy right-handed participants (10 females, 20-40, all with normal or corrected-to-normal vision) participated in the experiment. All participants gave written consent prior to their participation and received payment for their participation in the experiment. Imaging was performed according to safety guidelines approved by the ethics committee of the Institute for Research in Fundamental Sciences (IPM). Data from 2 participants was removed from the analysis because of excessive head motion (more than 3 mm, throughout the session).

### Stimuli and experimental procedure

Stimuli were from the two categories of human bodies and houses similar to the ones used in previous studies ^26,27,28^. Each category consisted of ten unique exemplars in gray-scale format (Figure 1a). These exemplars differed in identity, pose (for bodies), and viewing angle (for houses). Stimuli were fitted into a transparent square subtending 10.2°of visual angle and placed on a grey background. A central red fixation point subtending 0.45°of visual angle was present throughout the run. Stimuli from each category were made half-transparent and were presented either in isolation or superimposed on stimuli from the other category.

In a blocked-design paradigm, participants were instructed by a word cue to attend to bodies, houses, or the color of the fixation point. In blocks with attention to bodies or houses, participants performed a one-back repetition detection task on the attended stimuli by pressing a button when the exact same stimulus appeared in two consecutive trials. In blocks with attention directed towards the fixation point color, participants responded when the color of the fixation point changed from red to orange. These blocks served as conditions in which the visual stimuli (bodies and houses) were ignored. During each block, ten exemplars from the same category were presented either in isolation or superimposed on ten exemplars from the other stimulus category. Each image was presented for 400 ms, followed by 400 ms of fixation. Repetition occurred 2-3 times at random in each block. Each run consisted of 7 blocks with all possible combinations of stimulus categories and attention: Attend isolated bodies, Attend cluttered bodies, Attend isolated houses, Attend cluttered houses, Ignore isolated bodies, Ignore isolated houses, Ignore cluttered bodies and houses. The presentation order of the blocks was counterbalanced across the experimental runs. Each run lasted 2 min 14 s. For the main experiment, 14 participants completed 16 runs, 2 participants completed 14 runs, and 3 participants completed 12 runs.

### Localizer Experiments

In this study we examined the primary visual cortex (V1) along with the object-selective areas lateral occipital cortex (LOC) and posterior fusiform (pFs), the body-selective extrastriate body area (EBA), and the scene-selective parahippocampal place area (PPA). All participants completed four localizer runs which were used to define the primary visual and the category-selective regions of interest (ROIs). We used previously established protocols for the localizer experiments, but the details are repeated here for clarification and convenience.

#### Early Visual Area Localizer

We used meridian mapping to localize the primary visual cortex V1. Participants viewed a black-and-white checkerboard pattern through a 60 degree polar angle wedge aperture. The wedge was presented either horizontally or vertically. Participants were asked to detect a luminance changes in the wedge in a blocked-design paradigm. Each run consisted of four horizontal and four vertical blocks, each lasting 16 s, with 16 s of fixation in between. A final 16 s fixation followed the last block. Each run lasted 272 s. The order of the blocks was counterbalanced within each run. Each participant completed two runs of this localizer.

#### Category Localizer

A category localizer was used to localize the cortical regions selective to scenes, bodies, and objects. In a blocked-design paradigm, participants viewed stimuli from the five categories of faces, scenes, objects, bodies, and scrambled images. Each localizer run contained two 16-s blocks of each category, with the presentation order counterbalanced within each run. An 8-s fixation period was presented at the beginning, middle, and end of the run. In each block, 20 stimuli from the same category were presented. Each trial lasted 750 ms with 50 ms fixation in between. Participants were asked to maintain their fixation on a red circle at the center of the screen and press a key when they detected a slight jitter in the stimuli. Participants completed two runs of this localizer, each lasting 344 s. LOC ^29^, pFs ^30^ EBA ^31^ and PPA ^32^ were then defined using this category localizer.

### Data acquisition

Data were collected on a Siemens Prisma MRI system using a 64-channel head coil at the National Brain-mapping Laboratory (NBML). For each participant, we performed a whole-brain anatomical scan using a T1-weighted MP-RAGE sequence at the beginning of data acquisition. For the functional scans, including the main experiment and the localizer experiments, 33 slices parallel to the AC-PC line were acquired using T2*-weighted gradient-echo echo-planar sequences covering the whole brain(TR=2 s, TE=30 ms, flip angle = 90°, voxel size=3 × 3 × 3 mm^3^, *matrixsize*= 64 × 64). The stimuli were back-projected onto a screen using an LCD projector with the refresh rate of 60 Hz and the spatial resolution of 768 × 1024 positioned at the rear of the magnet, and participants observed the screen through a mirror attached to the head coil. MATLAB and Psychtoolbox were used to create all stimuli.

### Preprocessing

fMRI data analysis was performed using FreeSurfer (https://surfer.nmr.mgh.harvard.edu) and in-house MATLAB codes. fMRI data preprocessing steps included 3D motion correction, slice timing correction, and linear and quadratic trend removal. No spatial smoothing was performed on the data. A double gamma function was used to model the hemodynamic response function. We eliminated the first four volumes of each run to allow the signal to reach a steady state.

### Region of interest (ROI) definition

The primary visual cortex V1 was determined using a contrast of horizontal versus vertical polar angle wedges to reveal the topographic maps in the occipital cortex. ^33,34^. To define the object-selective areas LOC in the lateral occipital cortex and pFs in the posterior fusiform gyrus ^29,30^, we used a contrast of objects versus scrambled images. Clusters of active voxels in the lateral occipital and ventral occipitotemporal cortex were selected as LOC and pFS, respectively, following the procedure described by Kourtzi and Kanwisher ^35^. We used a contrast of scenes versus objects for defining the scene-selective area PPA in the parahippocampal gyrus ^32^, and a contrast of bodies versus objects for defining the body-selective area EBA in the lateral occipitotemporal cortex ^31^. The activation maps for both localizers were thresholded at *p* < 0.001, uncorrected.

### Data analysis

We performed a general linear model (GLM) analysis for each participant to estimate voxel-wise coefficients for each task condition. We then used these coefficients to compare the BOLD response in different conditions in each ROI. Preferred and null categories for each voxel were determined using voxel responses in conditions with isolated stimuli with the participant performing the task on the fixation point (color blocks). We then determined the activity in seven conditions for each voxel: P^at^, P^at^N, PN^at^, N^at^, P, PN, N, with P and N denoting the presence of the preferred and null and stimuli and the superscript at denoting the attended category. For instance, P refers to the condition in which the unattended preferred stimulus was presented in isolation, and P^at^N refers to the paired condition with the attended preferred and unattended null stimuli.

We used three models to predict the results: the weighted sum, weighted average and normalization models. The weighted sum model is a simple linear model suggesting that the response to multiple stimuli is the sum of responses to individual stimuli, and attention to a stimulus increases the response to that stimulus by the attention-related parameter, β:

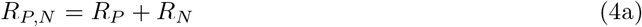

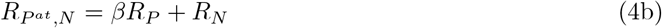

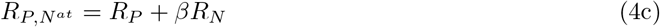

with R_P, N_ denoting the response elicited with the preferred and null stimuli present in the receptive field, and R_P_ and R_N_ denoting the response to isolated preferred and null stimuli, respectively. The superscript *at* specifies the attended stimulus, and the stimulus is ignored otherwise.

The weighted average model ^10,11,23^ is also a linear model that posits the response to multiple stimuli as the average of individual responses. Similar to the weighted sum model, the response of an attended stimulus is enhanced by the parameter related to attention, β:

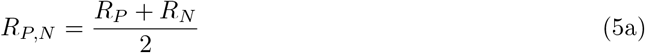

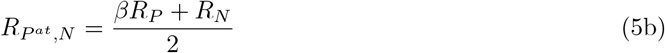

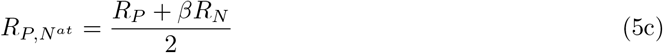

The normalization model of attention ^1,2,5,7^ can be described using divisive normalization with a saturation term in the denominator:

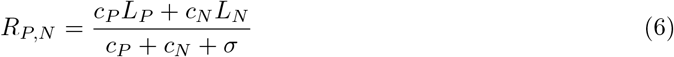

Here, L_P_and L_N_ denote the excitatory drive induced by the preferred or the null stimulus, respectively and σ represents the semi-saturation constant. c_P_ and c_N_ are the respective contrasts of the stimuli. Zero contrast denotes that the respective stimulus is not present in the visual field. When attention is directed towards one of the stimuli, we can rewrite Equation 6 as:

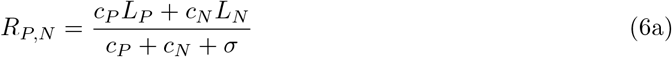

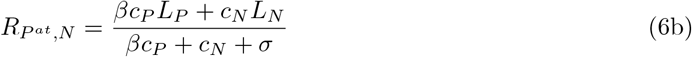

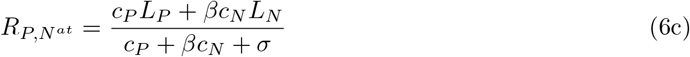

We fit model parameters for the three models. β was fit as a free parameter for all models. The normalization model had three additional free parameters, *L_p_*, *L_n_*, and *σ*. σ and β were constrained to be greater than zero and one, respectively, and less than 10. *L_p_* and *L_n_* were constrained to have an absolute value of less than 10. We estimated model parameters using constrained nonlinear optimizing, which minimized the sum-of-square errors. We split the fMRI data into two halves of odd and even runs and estimated model parameters for the first half as described. Then, using the estimated parameters for the first half, we calculated model predictions for each voxel in each condition and compared the predictions with the left-out half of the data. All comparisons of data with models, including the calculation of the goodness of fit, were done using the left-out data. Since the weighted sum and weighted average models used the response in the P and N conditions to predict responses in the remaining five conditions, we only used these five conditions and excluded the P and N conditions when calculating the goodness of fit for all models. The goodness of fit was calculated by taking the square of the correlation coefficient between the observed and predicted responses for each voxel across the five modeled conditions. We also calculated the correlation between voxel responses of the two halves of data across the same five conditions and calculated the noise ceiling in each ROI as the squared coefficient of this correlation. We defined two indices to quantify the observed effects of attention. The first index was used to compare voxel activities in paired conditions in which attention was directed toward the objects. We defined the response change index as the difference in average voxel activity when attention shifted from the preferred to the null stimulus, (*P^at^N* – *PN^at^*). The second index was used to quantify the asymmetry in attentional modulation. The asymmetry index, [(*PN^at^* – *N^at^*) – (*P^at^* – *P^at^N*)], compared the effect of the unattended stimulus on the response in conditions with unattended preferred or null stimuli. The comparison of observed indices with indices calculated from model predictions were done using the left-out part of the data.

## Acknowledgement

Maryam Vaziri-Pashkam was supported by NIH Intramural Research Program ZIA-MH002035.

## Author contributions

M.V.P. and N.D. designed the research. N.D. performed the research analyzed the data and wrote the paper. M.V.P. and G.H.Z. supervised the project at all stages and contributed to the writing.

## References

[1] David J Heeger. Normalization of cell responses in cat striate cortex. Visual neuroscience, 9(2):181–197, 1992.

[2] Matteo Carandini, David J Heeger, and J Anthony Movshon. Linearity and normalization in simple cells of the macaque primary visual cortex. Journal of Neuroscience, 17(21):8621–8644, 1997.

[3] Matteo Carandini and David J Heeger. Normalization as a canonical neural computation. Nature Reviews Neuroscience, 13(1):51–62, 2012.

[4] Pinglei Bao and Doris Y Tsao. Representation of multiple objects in macaque category-selective areas. Nature communications, 9(1):1–16, 2018.

[5] John H Reynolds and David J Heeger. The normalization model of attention. Neuron, 61(2):168–185, 2009.

[6] Joonyeol Lee and John HR Maunsell. Attentional modulation of mt neurons with single or multiple stimuli in their receptive fields. Journal of Neuroscience, 30(8):3058–3066, 2010.

[7] Amy M. Ni, Supratim Ray, and John H.R. Maunsell. Tuned normalization explains the size of attention modulations. Neuron, 73(4):803–813, 2012.

[8] Amy M Ni and John HR Maunsell. Neuronal effects of spatial and feature attention differ due to normalization. Journal of Neuroscience, 39(28):5493–5505, 2019.

[9] Ilona M Bloem and Sam Ling. Normalization governs attentional modulation within human visual cortex. Nature communications, 10(1):1–10, 2019.

[10] Davide Zoccolan, David D Cox, and James J DiCarlo. Multiple object response normalization in monkey inferotemporal cortex. Journal of Neuroscience, 25(36):8150–8164, 2005.

[11] Sean P MacEvoy and Russell A Epstein. Decoding the representation of multiple simultaneous objects in human occipitotemporal cortex. Current Biology, 19(11):943–947, 2009.

[12] Leila Reddy, Nancy G Kanwisher, and Rufin VanRullen. Attention and biased competition in multi-voxel object representations. Proceedings of the National Academy of Sciences, 106(50):21447–21452, 2009.

[13] Libi Kliger and Galit Yovel. The functional organization of high-level visual cortex determines the representation of complex visual stimuli. Journal of Neuroscience, 40(39):7545–7558, 2020.

[14] Jeffrey Moran and Robert Desimone. Selective attention gates visual processing in the extrastriate cortex. Science, 229(4715):782–784, 1985.

[15] Robert Desimone and John Duncan. Neural mechanisms of selective visual attention. Annual review of neuroscience, 18(1):193–222, 1995.

[16] Marvin M Chun, Julie D Golomb, and Nicholas B Turk-Browne. A taxonomy of external and internal attention. Annual review of psychology, 62:73–101, 2011.

[17] Stefan Treue and Julio C Martinez Trujillo. Feature-based attention influences motion processing gain in macaque visual cortex. Nature, 399(6736):575–579, 1999.

[18] Carrie J McAdams and John HR Maunsell. Effects of attention on orientation-tuning functions of single neurons in macaque cortical area v4. Journal of Neuroscience, 19(1):431–441, 1999.

[19] John H Reynolds, Tatiana Pasternak, and Robert Desimone. Attention increases sensitivity of v4 neurons. Neuron, 26(3):703–714, 2000.

[20] Julio C Martinez-Trujillo and Stefan Treue. Attentional modulation strength in cortical area mt depends on stimulus contrast. Neuron, 35(2):365–370, 2002.

[21] Amy M Ni and John HR Maunsell. Spatially tuned normalization explains attention modulation variance within neurons. Journal of neurophysiology, 118(3):1903–1913, 2017.

[22] Kendrick N Kay, Jonathan Winawer, Aviv Mezer, and Brian A Wandell. Compressive spatial summation in human visual cortex. Journal of neurophysiology, 110(2):481–494, 2013.

[23] Annelies Baeck, Johan Wagemans, and Hans P Op de Beeck. The distributed representation of random and meaningful object pairs in human occipitotemporal cortex: the weighted average as a general rule. Neuroimage, 70:37–47, 2013.

[24] Erik P Cook and John HR Maunsell. Attentional modulation of behavioral performance and neuronal responses in middle temporal and ventral intraparietal areas of macaque monkey. Journal of Neuroscience, 22(5):1994–2004, 2002.

[25] Laura Busse, Alex R Wade, and Matteo Carandini. Representation of concurrent stimuli by population activity in visual cortex. Neuron, 64(6):931–942, 2009.

[26] Maryam Vaziri-Pashkam and Yaoda Xu. Goal-directed visual processing differentially impacts human ventral and dorsal visual representations. Journal of Neuroscience, 37(36):8767–8782, 2017.

[27] Maryam Vaziri-Pashkam and Yaoda Xu. An information-driven 2-pathway characterization of occipitotemporal and posterior parietal visual object representations. Cerebral Cortex, 29(5):2034–2050, 2019.

[28] Yaoda Xu and Maryam Vaziri-Pashkam. Task modulation of the 2-pathway characterization of occipitotemporal and posterior parietal visual object representations. Neuropsychologia, 132:107140, 2019.

[29] Rafael Malach, JB Reppas, RR Benson, KK Kwong, H Jiang, WA Kennedy, PJ Ledden, TJ Brady, BR Rosen, and RB Tootell. Object-related activity revealed by functional magnetic resonance imaging in human occipital cortex. Proceedings of the National Academy of Sciences, 92(18):8135–8139, 1995.

[30] Kalanit Grill-Spector, Tammar Kushnir, Talma Hendler, Shimon Edelman, Yacov Itzchak, and Rafael Malach. A sequence of object-processing stages revealed by fmri in the human occipital lobe. Human brain mapping, 6(4):316–328, 1998.

[31] Paul E Downing, Yuhong Jiang, Miles Shuman, and Nancy Kanwisher. A cortical area selective for visual processing of the human body. Science, 293(5539):2470–2473, 2001.

[32] Russell Epstein, Alison Harris, Damian Stanley, and Nancy Kanwisher. The parahippocampal place area: recognition, navigation, or encoding? Neuron, 23(1):115–125, 1999.

[33] Martin I Sereno, AM Dale, JB Reppas, KK Kwong, JW Belliveau, TJ Brady, BR Rosen, and RB Tootell. Borders of multiple visual areas in humans revealed by functional magnetic resonance imaging. Science, 268(5212):889–893, 1995.

[34] Roger BH Tootell, Nouchine K Hadjikhani, Wim Vanduffel, Arthur K Liu, Janine D Mendola, Martin I Sereno, and Anders M Dale. Functional analysis of primary visual cortex (v1) in humans. Proceedings of the National Academy of Sciences, 95(3):811–817, 1998.

[35] Zoe Kourtzi and Nancy Kanwisher. Cortical regions involved in perceiving object shape. Journal of Neuroscience, 20(9):3310–3318, 2000.

